# GatorDuo: Global-Consistency Dual-Graph Refinement With Pseudo-Label Agreement for Spatial Transcriptomics

**DOI:** 10.64898/2026.05.10.724039

**Authors:** Zhenhao Zhang, Antonio Jimeno Yepes, Jiang Bian, Fuyi Li, Yuxi Liu

## Abstract

Spatial transcriptomics (ST) measures gene expression together with spatial coordinates, enabling spatial domain identification of coherent tissue regions. Many recent approaches rely on graph-based modeling to combine spatial neighborhoods and transcriptomic (gene-expression) similarity, yet neighborhood construction is often unreliable under sparsity and technical noise. As a result, spurious cross-domain shortcut edges can persist in static graphs and propagate misleading signals during message passing, ultimately blurring domain boundaries and weakening cluster separability. In this paper, we propose GatorDuo, a topology-aware dual-graph contrastive self-supervised framework for robust spatial domain identification that couples gene-expression similarity with spatial proximity through complementary neighborhood graphs. GatorDuo introduces global-consistency–based graph refinement that uses a pseudo-label agreement mask to suppress cross-domain shortcut edges in both views, thus stabilizing neighborhood topology for representation learning. To avoid manual tuning of domain resolution, GatorDuo further employs a contextual bandit reinforcement-learning strategy to adaptively select the clustering granularity (the number of clusters) used for refinement. The refined view-specific embeddings are integrated via a hybrid-routing Mixture-of-Experts (MoE) module to generate a unified embedding, optimized with contrastive objectives augmented by an MoE-alignment term. Across eight public benchmarks spanning sequencing- and imaging-based ST at spot and single-cell resolution, and compared with ten representative baselines, GatorDuo consistently delivers strong and robust spatial domain identification performance across multiple clustering metrics, while yielding informative unified embeddings that can support downstream biological analyses.

## I. Introduction

Spatial transcriptomics (ST) profiles gene expression while preserving the spatial coordinates of each measured unit (e.g., a capture spot or a segmented cell) within intact tissue sections. By linking molecular states with their native microenvironment, ST has become increasingly relevant for translational research. In oncology, it can delineate tumor-associated cell-type interactions and microenvironmental organization, as demonstrated in HER2-positive breast cancer [1]. In neuroscience, spatial profiling supports the mapping of transcriptional programs at the atlas-scale to anatomical structures, exemplified by a molecular atlas of the adult mouse brain [2]. In addition to sequencing-based assays, imaging-based platforms enable highly multiplexed RNA measurements directly in tissue and at cellular resolution [3], [4]. These advances have also enabled spatiotemporal atlasing in developmental systems, including mouse organogenesis [5]. In these biomedical settings, a central computational objective is the identification of spatial domains, i.e., biologically coherent tissue regions that correspond to different cellular ecosystems and microanatomical organization [6].

From a computational modeling perspective, accurate domain identification typically relies on the integration of complementary sources of structure present in ST data. Spatial domains are often informed by two complementary sources of structure: local anatomical continuity among spatial neighbors [7] and nonlocal transcriptomic similarity among functionally related units that can be spatially separated [8]. Nonlocal transcriptomic similarity can complement spatial proximity for domain identification by linking recurrent molecular programs across distant regions, denoising sparse ST profiles via information sharing among expression-matched units, and mitigating cases where spatial proximity does not reflect molecular similarity (e.g., boundaries or mixed spots).

Despite these motivations, many existing approaches induce neighborhoods primarily by spatial proximity (or array geometry). They then treat gene expression mainly as node features that are to be propagated, smoothed, or embedded over a fixed spatial topology such that domain identification relies predominantly on coordinate-defined neighborhoods. For instance, BayesSpace [9] defines neighbors according to the capture geometry and promotes spatial coherence by encouraging adjacent spots to share cluster labels via a spatial prior. NichePCA [10] builds a kNN graph in coordinate space and aggregates the expression of each unit with that of its spatial neighbors before PCA and clustering. Similarly, GraphST, SpaGCN, and SpaceFlow construct graphs from spatial adjacency and learn representations by propagating transcriptomic features over these coordinate-defined neighborhoods [11]– [13]. Finally, spaVAE [14] does not explicitly construct a transcriptomic-similarity graph; instead, it encodes inter-spot dependence through a Gaussian-process prior parameterized by Euclidean distances between spatial coordinates. As a result, long-range relationships defined by the transcriptome are not explicitly instantiated during neighborhood construction in these methods, which can fragment recurring biological states into disconnected regions or yield anatomically implausible partitions in tissues where similar programs recur at distant locations.

To incorporate long-range similarities derived from the transcriptome, only a few approaches explicitly model coordinate- and expression-derived relations via dual graphs, i.e., two complementary graphs defined on the same set of spots/cells, where one encodes spatial proximity and the other links transcriptionally similar (potentially distant) units (including Spatial-MGCN [15], STMGCN [16], MuCoST [8], DMGCN [17], STMIGCL [18]). Nevertheless, the graph topology is treated as static once it is constructed. Under the sparsity and technical noise of ST measurements, these static graphs can retain noise-induced cross-domain shortcut edges, which are then amplified by message passing to mix signals across latent groups, blur domain boundaries, and reduce embedding separability, ultimately hindering accurate domain identification.

In this paper, we propose **GatorDuo**, a topology-aware dual-graph contrastive self-supervised framework for robust spatial domain identification, as shown in Figure 1. Specifically, **GatorDuo** (i) induces complementary expression- and coordinate-derived graphs, (ii) performs global-consistency– based topology refinement using a pseudo-label agreement mask to prune cross-domain shortcut edges in both views and stabilize neighborhoods, (iii) automatically selects the pseudo-label granularity via a contextual-bandit policy to match dataset-specific domain resolution, and (iv) integrates refined dual-graph representations using a hybrid-routing Mixture-of-Experts (MoE) module trained with contrastive objectives (including an MoE-alignment term) to obtain a unified, dis-criminative embedding.

**Fig. 1.**
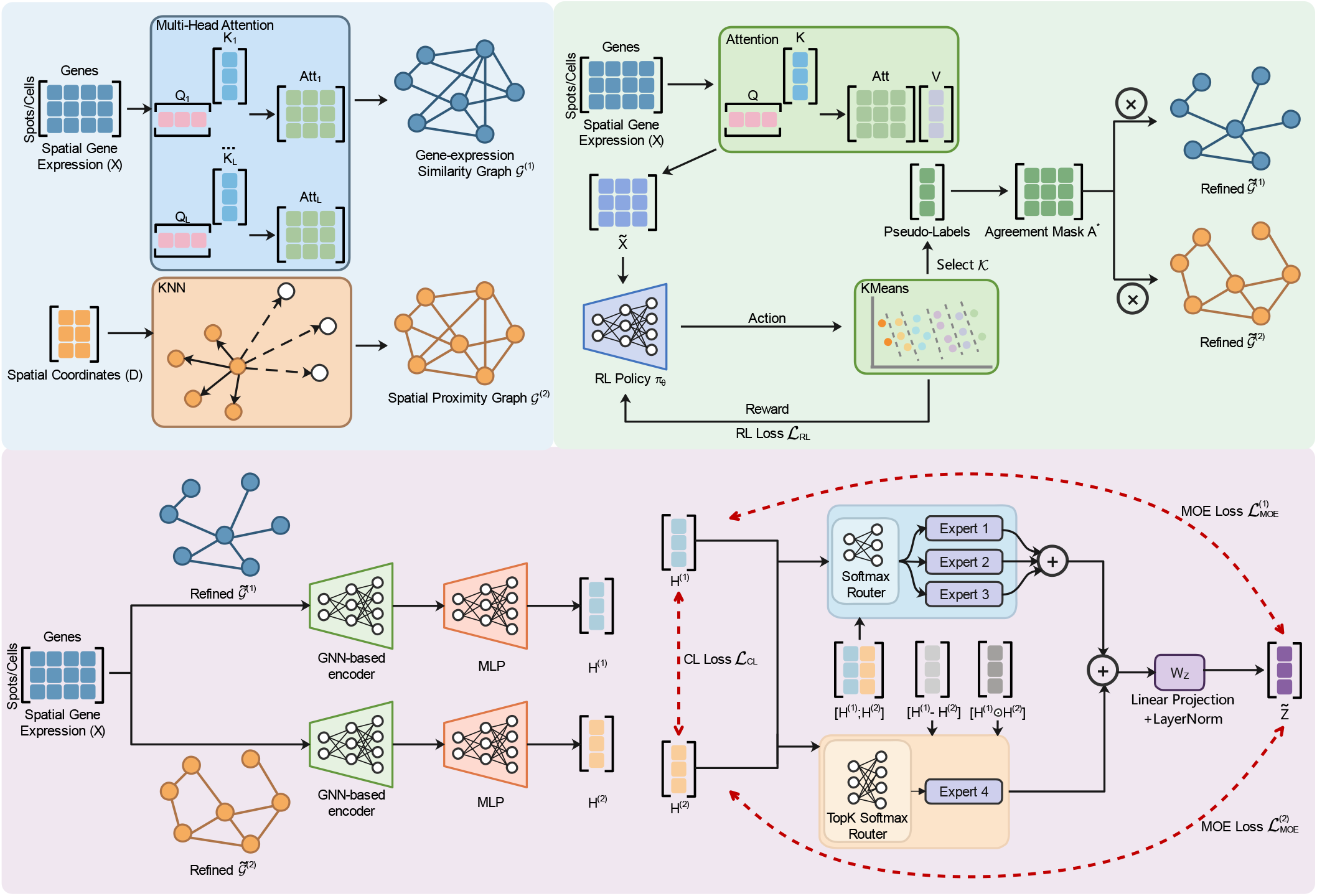
Overview of the proposed GatorDuo framework. Starting from X and D, we construct a dual-view graph representation that comprises a transcriptomic/gene-expression similarity graph 𝒢^(1)^ (via attention-derived Â and thresholded adjacency A^(1)^) and a spatial proximity graph 𝒢^(2)^ (via k-nearest-neighbor (kNN) adjacency A^(2)^). We obtain robust pseudo-labels by clustering attention-refined attributes 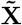 with an automatically selected 𝒦, and prune cross-domain (cross-cluster) shortcut edges in both views using the agreement mask A^*^, yielding refined adjacencies Ã^(1)^ and Ã^(2)^. The refined graphs are encoded into view-specific embeddings H^(1)^ and H^(2)^, fused into 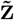 via a hybrid-routing Mixture-of-Experts module, and optimized by a unified objective that combines ℒ_*CL*_, ℒ_*MOE*_, and ℒ_*RL*_.

Our contributions are five-fold as follows:

- We propose **GatorDuo**, a topology-aware dual-graph contrastive self-supervised framework for robust spatial domain identification, coupling transcriptomic-similarity and spatial-proximity graphs within a unified iterative refinement framework.
- We introduce a **global-consistency topology refinement** scheme that derives reliable pseudo-labels (via attention-based attribute refinement) and prunes cross-domain shortcut edges in both views using a pseudo-label agreement mask, thereby stabilizing message-passing neighborhoods.
- We propose a **contextual-bandit reinforcement learning** strategy to automatically select the pseudo-label granularity used for topology refinement, enabling dataset-adaptive domain resolution without manual tuning.
- We design a **hybrid-routing MoE fusion** module trained with contrastive objectives (including an MoE-alignment term) to integrate refined dual-graph representations into a unified, discriminative embedding.
- Extensive experiments on **eight public benchmarks** across sequencing-based and imaging-based spatial profiling (spot- and cell-resolved), together with comparisons to **ten representative baselines**, ablations, and hyperparameter sensitivity analyses, demonstrate the effectiveness, cross-platform generalizability, and robustness of **GatorDuo**.

## II. Methods

### A. Dual-Graph Neighborhood Induction for ST Data

ST profiles each spot/cell with (i) a high dimensional gene expression vector and (ii) its physical coordinates on the tissue slide. Effective representation learning for ST data should jointly capture two complementary relationships: (i) spots/cells with similar expression profiles may share underlying biological states even if they are not spatially adjacent; and (ii) spatially proximal spots/cells often show local continuity due to tissue architecture and neighborhood effects. To this end, we represent each ST sample as two graphs defined over the same set of nodes: an expression-based graph that connects transcriptionally similar spots/cells, and a spatial graph that connects spatial neighbors.

#### a) Resolution and Node Definition

ST technologies can provide measurements at different resolutions. In spot-based platforms, a spot corresponds to a capture location and its expression profile may reflect transcripts from multiple cells, whereas in imaging-based platforms, a cell corresponds to an individual segmented cell. Throughout this paper, we adopt a unified abstraction and treat each spot or cell as a node with an expression profile and a spatial location; accordingly, both the expression-based graph and the spatial graph in our dual-graph neighborhood induction apply to either resolution.

Let 𝒢^(1)^ = (𝒱, ℰ^(1)^) and 𝒢^(2)^ = (𝒱, ℰ^(2)^) denote the gene-expression similarity graph and the spatial proximity graph, respectively. The two graphs share the same node set 𝒱 = {*v*_1_, *v*_2_, …, *v*_*N*_}, where *N* is the number of spots/cells. The edge sets ℰ^(1)^ and ℰ^(2)^ are induced by gene-expression similarity and spatial proximity, respectively. Let **X** ∈ ℝ^*N* ×*M*^ denote the gene expression matrix, where *M* is the number of genes. Let **D** ∈ ℝ^*N* ×2^ denote the spatial coordinates of spots/cells.

Based on the above definitions, we next describe how to construct the two graphs 𝒢^(1)^ and 𝒢^(2)^ from the observed gene expression matrix **X** and spatial coordinates **D**. Specifically, we derive the expression-based adjacency A^(1)^ by learning pairwise transcriptomic similarities, and build the spatial adjacency A^(2)^ by connecting proximal spots/cells in the coordinate space.

#### b) Dual-Graph Construction

To construct 𝒢^(1)^, we learn a pairwise similarity matrix from **X** using multi-head attention mechanism. Specifically, for each attention head (i.e., Head_*i*_), query and key representations are generated using learnable linear projections as follows:

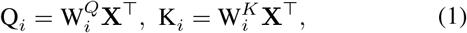

where all W are trainable parameters. We then compute the head-specific cell–cell (or spot–spot) similarity matrix by scaled dot-product attention as follows:

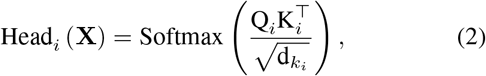

where 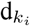 denotes the key/query dimension and softmax(·) is applied row-wise to produce normalized, non-negative attention weights, thereby producing normalized pairwise similarity scores. Each head is evaluated independently to capture complementary similarity patterns under different representation subspaces.

Accordingly, we aggregate all heads by concatenating the head-wise similarity matrices and applying a learnable linear transformation as follows:

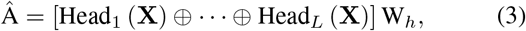

where ⊕ denotes concatenation, *L* is the number of heads, and W_*h*_ is a trainable parameter for fusing multi-head information. Here, Â serves as a dense similarity matrix, and we derive the final binary adjacency A^(1)^ by sparsifying Â via thresholding. In order to obtain a sparse adjacency matrix, we apply a learnable threshold *ψ* to Â as follows:

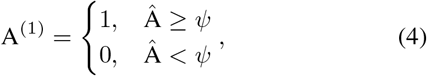

where *ψ* filters out weak associations and controls the sparsity of A^(1)^.

To construct 𝒢^(2)^, we build a k-nearest-neighbor (kNN) graph using spatial locations. For each pair of nodes (*v*_*i*_, *v*_*j*_), we compute their spatial distance *d*(*v*_*i*_, *v*_*j*_) from coordinates **D**. The hyperparameter *K*^(*KNN*)^ controls the neighborhood size (i.e., graph connectivity), with larger *K*^(*KNN*)^ yielding a denser local graph. The adjacency matrix A^(2)^ is defined by selecting up to *K*^(*KNN*)^ nearest neighbors for each node as follows:

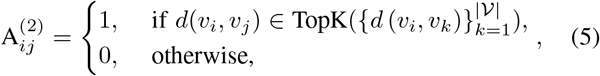

where TopK(·) corresponds to selecting the *K*^(*KNN*)^ nearest neighbors under the chosen distance-based criterion.

### B. Global-Consistency Graph Refinement with Pseudo-Label Agreement

The two initial graphs, 𝒢^(1)^ (expression-based) and 𝒢^(2)^ (spatial), are built by linking each node to a small set of neighbors (e.g., nearest neighbors in expression space). However, due to sparsity and technical noise in gene expression measurements and limited spatial sampling, local neighbor construction can be unreliable: two nodes may appear similar or spatially close by chance even though they belong to different spatial domains. As a result, the initial adjacencies may contain spurious cross-domain shortcut edges that connect nodes from different latent domains.

Such shortcut edges mix signals across domains during message passing, blurring domain boundaries and weakening the separability of the learned representations. To mitigate this issue, we introduce a clustering-induced global consistency prior by enforcing pseudo-label agreement at the edge level: node pairs assigned to different pseudo labels are regarded as inconsistent, and the corresponding edges are pruned. Accordingly, we construct a pseudo-label agreement mask and apply it to filter both adjacency matrices. We first obtain reliable pseudo labels via attentive attribute refinement, and then use an agreement mask to prune inconsistent edges in both graphs.

#### a) Attentive Attribute Refinement for Robust Pseudo-Labels

Because the pseudo-label agreement mask is induced by pseudo-label assignments, the quality of graph pruning depends on the fidelity of the pseudo labels. Directly clustering the raw attributes **X** can be sensitive to measurement noise, sparsity, and heterogeneous feature scaling, yielding noisy pseudo labels and consequently overly aggressive edge removal.

To improve clustering robustness, we employ an attention-based attribute enhancement module that performs data-adaptive attribute reweighting and aggregation, yielding an enhanced representation 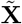. Specifically, we project **X** into query, key, and value spaces and compute 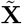 via scaled dot-product attention as follows:

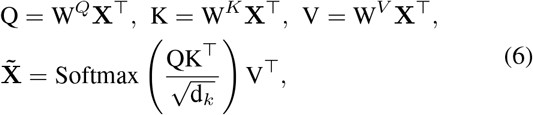

where all W are trainable parameters, Q, K and V denote the query, key, and value matrices, respectively; and d_*k*_ is the key dimension used for scaling. The output 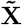 serves as the input to clustering, from which we derive pseudo labels for graph refinement.

Given 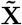, we apply KMeans to obtain pseudo-label assignments and cluster centers as follows:

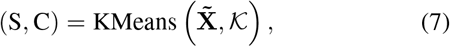

where S denotes the pseudo-label assignments and C denotes the corresponding cluster centers. 𝒦 is the number of clusters. Based on the obtained pseudo-labels, we build an agreement mask as follows:

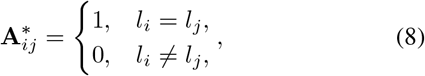

where *l*_*i*_ and *l*_*j*_ are the pseudo-labels of nodes *i* and *j*, respectively. We then refine the adjacency matrices of the two graphs via element-wise filtering (i.e., Hadamard product) as follows:

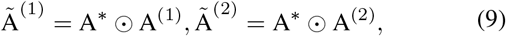

where A^*^ enforces that only pairs assigned to the same pseudo cluster can remain connected, thereby refining both 𝒢^(1)^ and 𝒢^(2)^ using clustering-consistent relations. Here, Ã^(1)^ and Ã^(2)^ are the refined adjacency matrices of 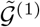 and 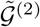, respectively.

#### b) Policy-Guided Auto-Tuning of Cluster Granularity

The pseudo-label agreement mask is constructed from KMeans-based pseudo labels on the enhanced representations, and its effectiveness therefore depends critically on the chosen number of clusters, i.e., 𝒦. The refinement is sensitive to 𝒦: overly small values may merge distinct tissue regions, whereas overly large values can fragment coherent domains and lead to unstable edge pruning. To avoid manual tuning and adaptively select an appropriate 𝒦 across datasets and training stages, we learn a data-dependent selector for 𝒦. Since the quality of a candidate 𝒦 can only be assessed after clustering, we treat 𝒦 selection as a contextual bandit problem and optimize a stochastic policy using immediate clustering-quality feedback.

Reinforcement learning provides a framework for learning decision policies from scalar feedback through interaction with an environment [19]. An RL agent interacts with an environment in a trial-and-error manner: at each decision step, it observes a state *s* that summarizes the current context, selects an action *a*, and receives a scalar reward *r* that reflects the quality of the resulting outcome. The agent then updates its policy to maximize the expected reward. In our formulation, the agent corresponds to the data-dependent selector. The state *s* is constructed from the current node representations, an action corresponds to choosing a candidate cluster number 𝒦, and the reward *r* is computed based on the resulting clustering quality.

Although the choice of *K* influences subsequent training through updated pseudo labels and graph structure, explicitly modeling long-horizon dependencies would substantially complicate optimization. From a modeling perspective, the resulting learning-and-clustering loop can be interpreted as a non-stationary Markov decision process (MDP) [20]. However, because *K* is selected once per iteration and its effect is evaluated immediately after the KMeans step, we adopt a contextual bandit (one-step) formulation and optimize each decision using only its immediate reward [21]. This choice yields a simple and stable training procedure that aligns with the per-iteration, one-step nature of KMeans clustering. We parameterize the selector as a stochastic categorical policy *π*_*θ*_(*a* | *s*) and optimize it with REINFORCE [22], [23], using the immediate clustering-quality reward to update *θ*. As a result, the selector adaptively chooses the cluster granularity without manual tuning.

To instantiate the above contextual bandit formulation, at each training step *t*, we specify the state representation, action space, policy parameterization, reward definition, and optimization objective.

#### c) State representation

The policy requires a fixed-dimensional input that does not depend on the selected cluster number 𝒦; otherwise, changing 𝒦 would change the input structure and destabilize training. Therefore, at training step *t*, we first obtain the enhanced node representations 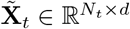 under the current model parameters, and then summarize them into a state vector **s**_*t*_ ∈ ℝ^2*d*^ using global mean and (element-wise) standard deviation as follows:

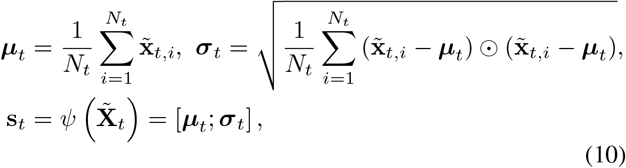

where 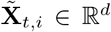 is the representation of node *i, N*_*t*_ is the number of nodes, and ⊙ denotes the element-wise product.

#### d) Action space

Given the state **s**_*t*_, the agent selects the number of clusters for the next KMeans step. Since 𝒦 is discrete and we restrict the search to a bounded interval [𝒦_min_, 𝒦_max_] for efficiency, we define a finite action set as follows:

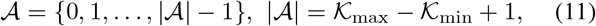

and map each action *a*_*t*_ ∈ 𝒜 to a cluster number by

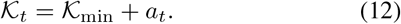

#### e) Policy parameterization

We model cluster-number selection as a categorical policy over 𝒜. At step *t*, the policy network *f*_*θ*_ (e.g., MLP) outputs logits **u**_*t*_ and induces a distribution via softmax as follows:

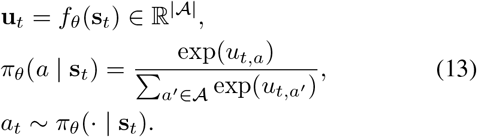

At inference, we use the greedy decision *a*_*t*_ = arg max_*a*∈𝒜_ *π*_*θ*_(*a* | **s**_*t*_) to obtain a deterministic 𝒦_*t*_.

#### f) Reward definition

After sampling *a*_*t*_ and obtaining 𝒦_*t*_, we run KMeans on 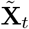 produce pseudo labels and cluster centers 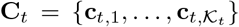 The pseudo labels are used for graph refinement (as described in the previous section), while the clustering result also provides a scalar reward to train the selector. Specifically, we define a clustering-oriented reward that encourages compact clusters and well-separated centers as follows:

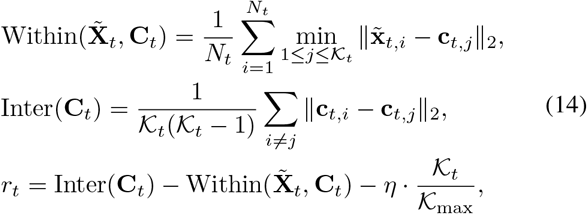

where *η* ≥ 0 penalizes overly large 𝒦_*t*_ to avoid unnecessary fragmentation.

#### g) Optimization objective

Under a contextual bandit formulation, we maximize the expected immediate reward as follows: 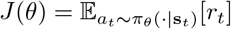. We optimize the policy using REINFORCE with entropy regularization as follows:

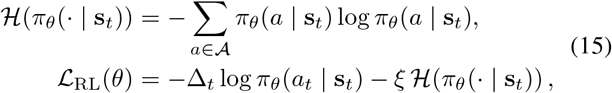

where *ξ >* 0 is the entropy coefficient that encourages exploration. To reduce the variance of the Monte Carlo policy-gradient estimator, we introduce an action-independent reward baseline *b*_*t*_ and define the advantage as Δ_*t*_ = *r*_*t*_ − *b*_*t*_. We implement *b*_*t*_ as an exponential moving average of past rewards as follows:

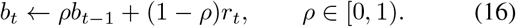

Subtracting this baseline does not change the expected policy gradient, but it typically yields more stable optimization by reducing gradient variance. With this design, cluster-number selection is trained jointly with unsupervised representation learning, enabling KMeans-based graph refinement without manually fixing the cluster number.

### C. Mixture-of-Experts Fusion with Hybrid Routing for Cross-Graph Integration

The refined expression and spatial graphs capture complementary neighborhood cues from transcriptomic and spatial perspectives. To obtain a single unified representation that exploits both cues, we design a Mixture-of-Experts (MoE) fusion module to adaptively integrate the two graph-specific representations. A MoE model consists of multiple expert subnetworks and a gating (routing) network that assigns each input to experts in a data-dependent manner [24]. In our implementation, the router outputs mixture coefficients over experts: we use a softmax router for the representation experts (dense mixture) and a TopKSoftmax (top-*k*) router for the interaction experts (sparse routing) [25], [26]. The fused output is computed as a weighted sum of expert outputs. This conditional computation increases model capacity while allowing different experts to specialize in distinct patterns, enabling adaptive fusion of heterogeneous signals across views.

Specifically, we encode each refined graph to obtain view-specific node embeddings, and then fuse them via expert routing. Given the two (refined) graphs 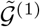 and 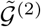, we compute the embeddings of the graph-specific nodes using a graph neural network (GNN) based encoder [27] followed by an MLP projection. Specifically, for the node feature (gene expression) matrix **X** and the adjacency matrices Ã^(1)^ and Ã^(2)^, we obtain the two view-specific representations as follows:

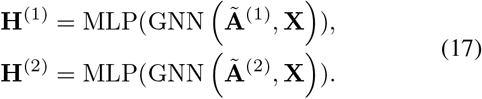

The MoE fusion module takes **H**^(1)^ and **H**^(2)^ as input and output a fused representation 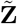. To make the fusion process more structured and controllable, we organize experts into two groups: representation experts and interaction experts. Representation experts operate on **H**^(1)^, **H**^(2)^, and [**H**^(1)^; **H**^(2)^] to capture view-specific information as well as cross-view consistent signals. In contrast, interaction experts take **H**^(1)^, **H**^(2)^, (**H**^(1)^ − **H**^(2)^), and (**H**^(1)^ ⊙ **H**^(2)^) as input to explicitly model feature-wise agreement, discrepancy, and co-activation patterns between the two representations. We next detail these two expert groups in turn.

#### a) Representation Experts

We instantiate representation experts for three inputs: **H**^(1)^, **H**^(2)^ and the concatenation [**H**^(1)^; **H**^(2)^]. Each expert is implemented as a two-layer MLP with GELU activation. For a generic input feature **u**, an expert can be written as follows:

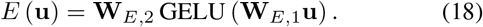

Accordingly, for the three inputs **H**^(1)^, **H**^(2)^ and [**H**^(1)^; **H**^(2)^], we compute the corresponding expert outputs as follows:

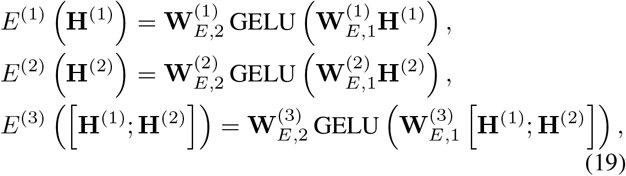

where all **W** are trainable parameters and GELU(·) is the GELU activation function.

#### b) Soft Routing (Router)

Each input branch is equipped with a linear router followed by a softmax to generate mixture weights as follows:

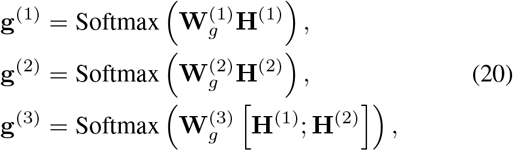

where all W are trainable parameters. Here, **g**(·) represents mixture coefficients over experts. The softmax is taken over the expert dimension so that each node receives a normalized set of routing weights.

#### c) Mixture Aggregation

The three intermediate representations are obtained via softmax-weighted aggregation as follows:

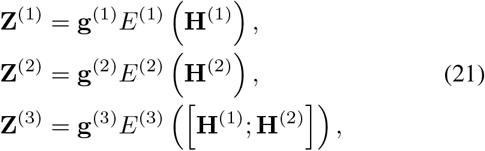

where the product **g***E*(·) denotes a weighted sum over expert outputs (i.e., a mixture over multiple MLP experts). The representation experts preserve (i) view-specific information via the separate mixtures on **H**^(1)^ or **H**^(2)^, and (ii) cross-view consistent information via the mixture on [**H**^(1)^; **H**^(2)^]. However, they do not explicitly encode feature-wise agreement or discrepancy between **H**^(1)^ and **H**^(2)^.

#### d) Interaction Experts

We introduce the interaction experts by constructing an interaction feature that concatenates the raw representations and simple element-wise interactions as follows:

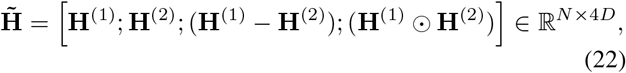

where ⊙ denotes the Hadamard product. The concatenated terms encode the original view-specific embeddings, their feature-wise discrepancy, and co-activation patterns, thereby providing an explicit basis for modeling cross-view correspondence. We then apply an MLP expert to 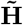 and use a TopKsoftmax router to obtain sparse routing weights as follows:

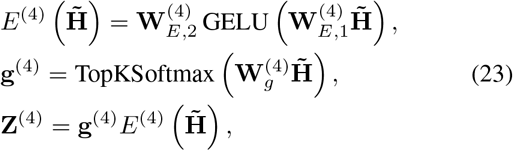

where TopKSoftmax(·) keeps only the top-k routing logits for each node, masks the remaining logits to −∞, and then applies softmax. This produces a sparse routing distribution, encouraging the interaction branch to concentrate on a small subset of experts for each node.

In the final fusion stage, we concatenate the four intermediate representations **Z**^(1)^, **Z**^(2)^, **Z**^(3)^, **Z**^(4)^ and apply a linear projection followed by LayerNorm to obtain the fused representation as follows:

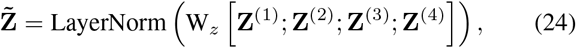

where W_*z*_ is a trainable parameter and 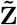 is the final output of the MoE fusion module.

### D. Contrastive Self-Supervision with MoE Alignment

After obtaining the fused representation 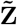, we train the model with an instance-discrimination contrastive self-supervised objective defined on the two view-specific representations generated by the two encoding branches, i.e., **H**^(1)^ and **H**^(2)^. Contrastive learning (CL) encourages view-consistent representations by pulling together embeddings of the same sample across views (positives) while pushing apart embeddings from different samples (negatives) [28]–[30]. We implement this objective using the InfoNCE cross-entropy loss within each mini-batch [31]. In particular, for each sample *i* in a mini-batch of *N*_*b*_, 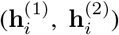 forms a positive pair, while the representations of the remaining samples in the mini-batch serve as negatives. We denote the two sets of mini-batch representations as follows:

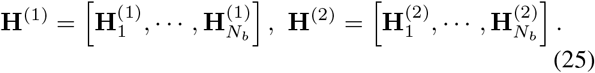

We then concatenate the two sets of representations as follows:

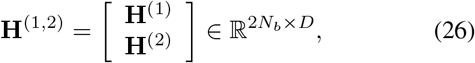

where 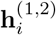 denotes the i-th row of **H**^(1,2)^. We treat 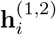 as an anchor, and define its positive index *p*(*i*) as follows:

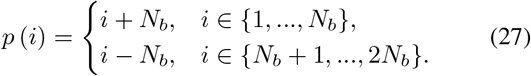

All remaining embeddings in the batch, 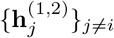, are treated as negatives for the anchor 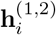. For each anchor 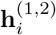, we minimize the cross-entropy loss that identifies its positive counterpart indexed by *p*(*i*) among all 2*N*_*b*_ candidates as follows:

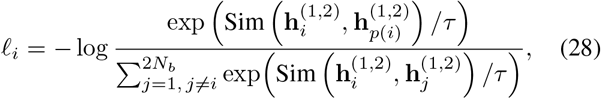

where Sim(·,·) denotes the similarity function, implemented as a cosine similarity, *τ* is the temperature hyperparameter. The final contrastive objective averages over all 2*N*_*b*_ anchors can be written as follows:

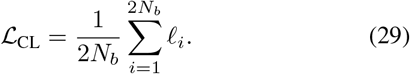

While ℒ_CL_ aligns the two branch embeddings **H**^(1)^ and **H**^(2)^ within an instance-discriminative space, it does not impose any constraint on the fused output 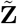, because ℒ_CL_ is computed solely from **H**^(1)^ and **H**^(2)^. Consequently, the MoE fusion module may produce fused embeddings that are not well aligned with the contrastive geometry learned by the encoders. To explicitly couple the fusion branch with the contrastive space, we introduce an MoE alignment loss as follows:

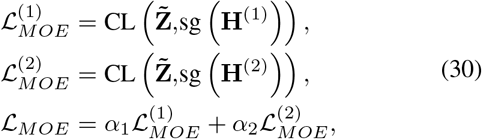

where CL(·,·) denotes the same contrastive objective defined above, and sg(·) is the stop-gradient operator. This design provides a direct self-supervised training signal for the MoE fusion branch: 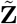 is optimized to match the instance-discriminative geometry induced by **H**^(1)^ and **H**^(2)^, while **H**^(1)^ and **H**^(2)^ act as fixed anchors under this term. The weights *α*_1_ and *α*_2_ control the relative contributions of the two reference representations.

Finally, we optimize the model with a unified objective that jointly accounts for (i) cross-view contrastive learning, (ii) MoE fusion alignment, and (iii) reinforcement learning reward used for adaptive cluster-number selection as follows:

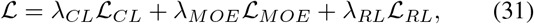

where *λ*_*CL*_, *λ*_*MOE*_ and *λ*_*RL*_ are trade-off coefficients that control the relative contributions of the three terms.

## III. Experiments

### A. Benchmark datasets and Baselines

We evaluated **GatorDuo** on a diverse suite of publicly available spatial transcriptomics datasets spanning multiple tissue types, biological contexts, and measurement technologies. For sequencing-based spatial transcriptomics, we included two widely used datasets: the HER2-positive breast tumor dataset [1] and the adult mouse brain dataset [2]. We further incorporated 10x Genomics Visium datasets from the 10x Genomics Data Repository, including one human breast cancer section^1^ and two mouse brain sagittal serial sections (anterior^2^ and posterior^3^). To further assess robustness and generalizability beyond Visium, we extended our benchmarking to imaging-based spatial molecular profiling datasets, MERFISH from the mouse hypothalamic preoptic region [3] and seqFISHPLUS from the mouse brain cortex [4] as well as a high-resolution sequencing-based platform, Stereo-seq, for atlasing mouse organogenesis [5]. Collectively, these datasets span broad ranges of spatial resolution, gene expression characteristics, and sampling density, providing a rigorous testbed for quantifying the generalizability, stability, and cross-platform robustness of the proposed **GatorDuo**.

We compared **GatorDuo** against a diverse set of established baselines for spatial representation learning and spatial domain identification in spatial transcriptomics, including Bayesian spatial modeling, graph neural networks, contrastive/self-supervised learning, and deep generative approaches. Because our evaluation spans both spot-level spatial transcriptomics datasets (ST-H1, ST-MAM, Human Breast Cancer, MOSTA, Mouse Brain Anterior (MB-Anterior), and Mouse Brain Posterior (MB-Posterior)) and cell-resolved imaging-based datasets (MERFISH and seqFISHPLUS), we grouped baselines according to their primary operating resolution: (i) methods tailored to spot-level measurements, (ii) methods designed for cell-resolved data, and (iii) resolution-agnostic methods that can be readily applied to either setting by treating each spatial unit (spot or cell) as an observation (e.g., SpaGCN and SpaceFlow; see descriptions below).

- **BayesSpace (spot-level)** [9] is a Bayesian spatial clustering framework that incorporates spatial neighborhood priors to improve spot-level domain detection and further supports enhanced-resolution inference at subspot granularity.
- **NichePCA (cell-level)** [10] is a simple and scalable PCA-based baseline for cell-resolved spatial domain identification: it constructs a cell neighborhood graph, aggregates local neighborhood expression into a “niche” representation, and applies PCA (followed by clustering) to identify spatial domains.
- **GraphST (spot-level)** [11] is a self-supervised graph representation learning method that builds a spot adjacency graph from spatial coordinates and uses contrastive learning to learn discriminative spot embeddings, supporting downstream tasks such as clustering, multi-sample integration, and scRNA-seq transfer/deconvolution onto spatial spots.
- **SpaGCN (spot/cell-level)** [12] is a graph convolutional framework that integrates gene expression with spatial coordinates (and optionally histology) by constructing a spatial graph over observations (spots or cells), enabling spatial domain detection and spatially variable gene identification.
- **SpaceFlow (spot/cell-level)** [13] is a deep graph model that learns spatially consistent low-dimensional embeddings by jointly encoding expression similarity and spatial proximity on a neighborhood graph, trained with a self-supervised/contrastive objective for downstream spatial domain identification.
- **spaVAE (spot-level)** [14] is a dependency-aware deep generative model that extends variational autoencoders with a hybrid latent prior (Gaussian process with Gaussian) to capture spatial correlations among observations, supporting tasks including clustering, denoising, spatial interpolation, and resolution enhancement.
- **Spatial-MGCN (spot-level)** [15] is a multi-view graph convolutional network that jointly models a feature graph and a spatial graph and uses attention-based fusion (with count-aware reconstruction) to derive embeddings for spatial domain identification.
- **MuCoST (spot-level)** [8] is a multi-view graph contrastive learning framework that fuses a spatial proximity view with an expression-similarity view to model both local adjacency and nonlocal co-expression dependencies, improving identification of spatial domains.
- **DMGCN (spot/cell-level)** [17] is a data-augmentation based multi-view graph convolutional framework for spatial domain identification that constructs a spatial neighbor graph and a gene-expression feature graph, and utilizes attention-based multi-view encoding with co-convolution and corrupted-feature contrastive learning to learn robust spots/cells embeddings, followed by a decoder to infer domain labels and reconstruct gene expression.
- **STMIGCL (spot-level)** [18] is a multi-view graph convolutional framework with implicit contrastive learning that builds multiple neighbor graphs from gene expression and spatial coordinates, refines low-dimensional representations via latent-space contrastive enhancement, and uses an attention mechanism to adaptively fuse multi-view embeddings for spatial domain identification and downstream analyses.

### B. Data pre-processing

We processed the raw count expression matrices from each spatial transcriptomics dataset using a standard preprocessing workflow to improve comparability across spots/cells and to stabilize variance for downstream modeling. Briefly, we first performed library-size normalization by dividing each gene’s count by the total counts within the corresponding spot/cell, thereby rescaling all spots/cells to a common sequencing depth. The normalized values were then multiplied by a fixed scale factor (default: 10^4^) and log-transformed to reduce the impact of extreme counts and improve numerical stability. A statistical summary of the datasets and samples analyzed in this study is provided in Table I.

**TABLE 1.**
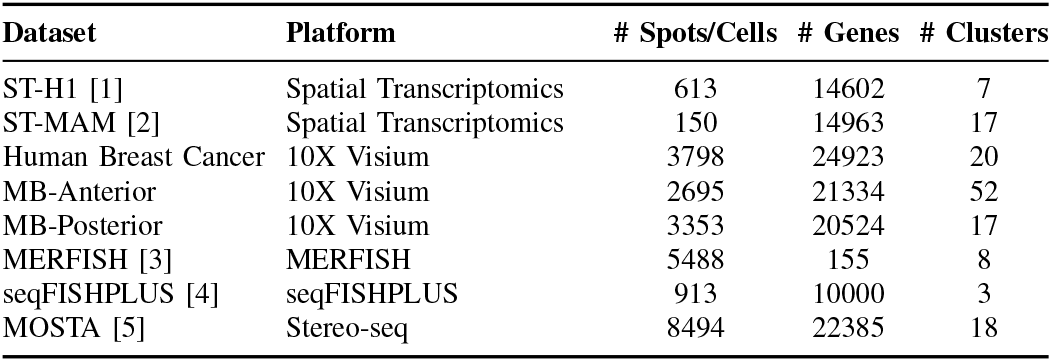
Summary of Benchmark Datasets, Platforms, and Statistics.

| Dataset | Platform | # Spots/Cells | # Genes | # Clusters |
| --- | --- | --- | --- | --- |
| ST-H1 [1] | Spatial Transcriptomics | 613 | 14602 | 7 |
| ST-MAM [2] | Spatial Transcriptomics | 150 | 14963 | 17 |
| Human Breast Cancer | 10X Visium | 3798 | 24923 | 20 |
| MB-Anterior | 10X Visium | 2695 | 21334 | 52 |
| MB-Posterior | 10X Visium | 3353 | 20524 | 17 |
| MERFISH [3] | MERFISH | 5488 | 155 | 8 |
| seqFISHPLUS [4] | seqFISHPLUS | 913 | 10000 | 3 |
| MOSTA [5] | Stereo-seq | 8494 | 22385 | 18 |

### C. Implementation and hyperparameter settings

The proposed **GatorDuo** was implemented in Python (v3.9.12) using PyTorch (v1.11.2). For each dataset, spots/cells were randomly split into training, validation, and test sets with an 80%/10%/10% ratio. For graph construction, we set the key/query dimension to 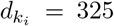 for each attention head and used *L* = 2 attention heads. The learnable threshold was initialized as *ψ* = 0.08, and the k-nearest-neighbor graph was built with *K*^(KNN)^ = 6. For global-consistency graph refinement, we set *d*_*k*_ = 282, 𝒦min = 3, 𝒦max = 20, *η* = 0.02, and *ξ* = 0.001. For MoE fusion, the hidden dimension of the GNN-based encoder was 217 and the MLP hidden dimension was 75, while the MoE dimension was set to 56. For contrastive self-supervision, the temperature hyperparameter was set to *τ* = 0.4. The overall objective was optimized as a weighted combination of multiple loss terms, with loss scaling coefficients *α*_1_ = 0.45 and *α*_2_ = 0.75, and *λ*_*CL*_ = 0.82, *λ*_*MOE*_ = 0.73, and *λ*_*RL*_ = 0.65. We trained the model using Adam with a learning rate of 3 × 10^−4^. The batch size was set to 52 for the MOSTA dataset, 95 for the MB-Anterior dataset, and 65 for all remaining datasets. Hyperparameters were selected via grid search, with validation performance used for model selection. For baseline methods, we used the default hyperparameters recommended by their respective packages. All experiments were conducted on a workstation equipped with an NVIDIA RTX 4090 GPU (24 GB memory).

### D. Evaluation metrics

To comprehensively evaluate spatial domain identification, we compared predicted clusters with ground-truth annotations using five widely adopted metrics: Adjusted Rand Index (ARI), Normalized Mutual Information (NMI), Clustering Accuracy (ACC), Purity, and Homogeneity. These metrics capture complementary aspects of agreement between predicted domains and reference labels, where higher scores indicate better performance. Specifically, ARI measures the agreement between the predicted and reference clusterings while accounting for randomness, and is therefore particularly useful when the class distributions are unbalanced or when clusters overlap. The ARI between the ground-truth labels **Y** and the predicted cluster assignments **Ŷ** can be written as follows:

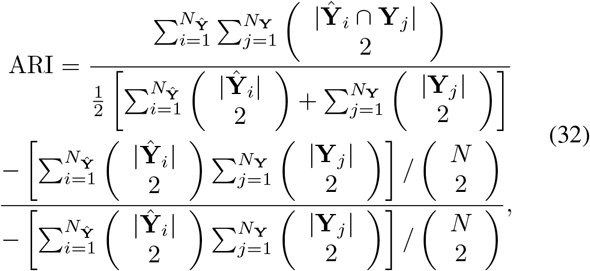

where *N*_**Y**_ and *N*_**Ŷ**_ denote the numbers of clusters in **Y** and **Ŷ**, respectively, and *N* is the total number of spots/cells.

The NMI quantifies the mutual dependence between the predicted and ground truth labels, while the normalization alleviates the influence of imbalanced label distributions. The NMI between **Y** and **Ŷ** can be written as follows:

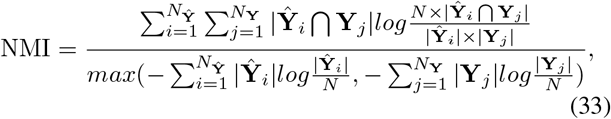

The ACC measures the proportion of correctly assigned cells after optimally matching predicted clusters to ground-truth labels. The ACC between **Y** and **Ŷ** can be written as follows:

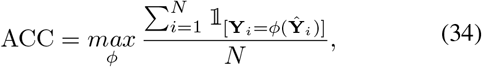

where *ϕ* is a mapping from predicted cluster indices to ground-truth labels, and 𝟙 _[·]_ denotes the indicator function.

The purity quantifies the extent to which each predicted cluster is dominated by a single ground-truth class, and is therefore commonly used to measure within-cluster label homogeneity. Intuitively, a clustering achieves high purity when most samples in each cluster share the same true label. Formally, given the ground-truth labels 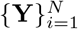 and predicted cluster assignments 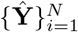, let *ϱ* = {*ϱ*_1_, *ϱ*_2_, · · ·, *ϱ*_|*ϱ*|_} denote the set of true classes and *ς* = {*ς*_1_, *ς*_2_,, · · ·, *ς*_|*ς*|_} denote the set of predicted clusters. For each cluster *ς*_*j*_, we identify the ground-truth class *ϱ*_*k*_ that appears most frequently within that cluster, and count those samples as correctly assigned. Purity is then computed as the fraction of correctly assigned samples over all *N* spots/cells:

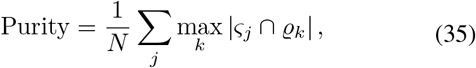

where |*ς*_*j*_ ∩ *ϱ*_*k*_| denotes the number of samples in cluster *ς*_*j*_ that belong to class *ϱ*_*k*_.

Finally, the homogeneity score evaluates the extent to which each cluster contains only cells from a single ground-truth class as follows:

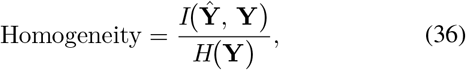

where *H*(**Y**) is the entropy of **Y** and *I*(**Ŷ**, **Y**) is the mutual information between **Ŷ** and **Y**. The score ranges from 0 to 1, with 1 indicating perfectly pure clusters (each cluster containing cells from only one class), whereas lower values indicate mixed clusters.

## IV. Results

### A. Spatial domain identification across diverse ST datasets

We evaluated **GatorDuo** on a diverse suite of publicly available spatial transcriptomics datasets spanning multiple tissue contexts and spatial resolutions. The benchmark covers sequencing-based spatial transcriptomics datasets (ST-H1 and ST-MAM), 10x Visium sections from human breast cancer and mouse brain sagittal slices, and high-resolution platforms including Stereo-seq MOSTA and single-cell–resolved imaging-based datasets (MERFISH and seqFISHPLUS). Notably, these benchmarks span a broad range of data regimes (as shown in Table I), from small sections with limited spatial units (e.g., 150–613 spots) to large-scale profiles with thousands of observations (up to 8,494 spots/cells), and from targeted gene panels (155 genes in MERFISH) to near whole-transcriptome measurements (over 20k genes in Visium/Stereo-seq). They also show different domain granularities (3–52 annotated clusters), reflecting substantial variation in underlying tissue complexity and annotation resolution.

We compared **GatorDuo** against representative Bayesian spatial modeling, graph neural networks, contrastive/self-supervised learning, and deep generative approaches, including BayesSpace, NichePCA, GraphST, SpaGCN, SpaceFlow, spaVAE, Spatial-MGCN, MuCoST, DMGCN, and STMIGCL. As summarized in Figure 2, **GatorDuo** achieves consistently strong performance for spatial domain identification across datasets under five widely used clustering metrics (ARI, NMI, ACC, Purity, and Homogeneity), which collectively quantify agreement between predicted domains and reference annotations. Across both spot-level and cell-resolved settings, **GatorDuo** consistently delivers competitive performance across datasets and metrics, indicating accurate recovery of tissue organization while maintaining high within-domain label consistency, as reflected by Purity and Homogeneity. Importantly, this strong performance is maintained across platforms with distinct characteristics (e.g., Visium versus MERFISH/seqFISHPLUS), supporting robust generalization. Overall, these results suggest that jointly modeling transcriptomic similarity and spatial proximity, together with graph-level consistency objectives, enhances the separability of latent tissue regions and enables robust spatial domain identification across diverse spatial transcriptomics benchmarks.

**Fig. 2.**
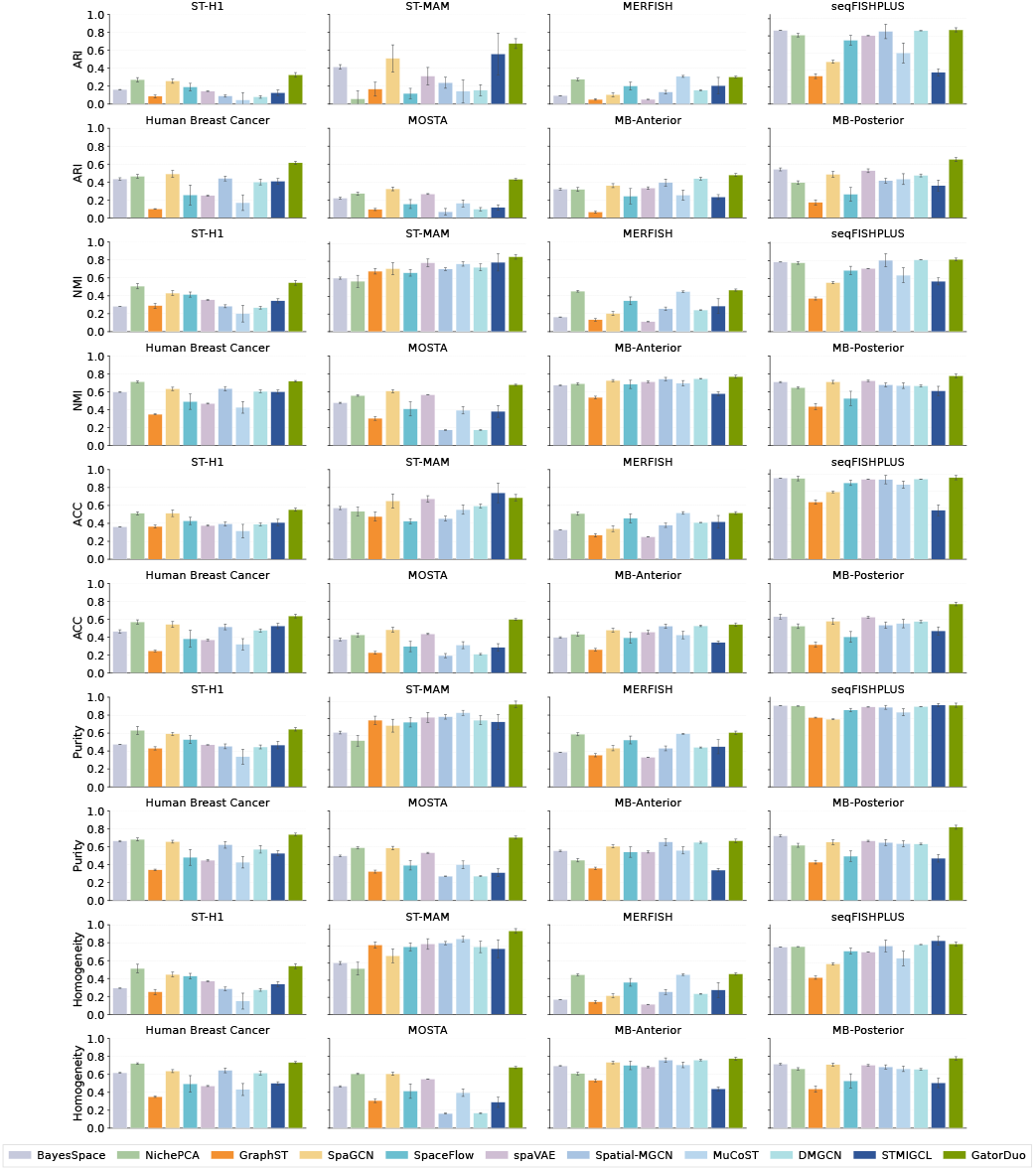
Spatial domain identification performance across eight spatial transcriptomics benchmarks. Performance comparison of GatorDuo with representative baselines on ST-H1, ST-MAM, Human Breast Cancer (Visium), MB-Anterior (Visium), MB-Posterior (Visium), MOSTA (Stereo-seq), MERFISH, and seqFISHPLUS. Results are evaluated using ARI, NMI, ACC, Purity, and Homogeneity; higher values indicate better agreement with reference annotations.

### B. Ablation study

To quantify the contribution of each component in **GatorDuo**, we conducted an ablation study with four variants. Specifically, **GatorDuo**_*α*_ (**w/o RL**) disables the RL-based selection of the cluster number *K* and instead uses a fixed *K* for pseudo-label generation. **GatorDuo**_*β*_ (**w/o update-graph**) removes the entire graph refinement module and keeps the initial graphs unchanged. **GatorDuo**_*γ*_ (**w/o MoE Loss**) retains the MoE fusion to obtain the combined representation but removes the MoE-specific loss term. **GatorDuo**_*δ*_ (**w/o MoE**) discards the MoE module and computes the combined embedding by simple averaging.

As shown in Figure 3, the full model consistently achieves the best overall performance across datasets and clustering metrics. Removing any individual component leads to a noticeable degradation, suggesting that these modules provide complementary benefits. Particularlly, omitting graph refinement (**GatorDuo**_*β*_) weakens the structural consistency of neighborhoods, resulting in less separable latent regions. This is consistent with our motivation that the initial graphs can contain noise-induced cross-cluster shortcut edges that mix information across latent groups and blur domain boundaries, whereas pseudo-label agreement pruning stabilizes the neighborhood topology. Replacing RL with a fixed *K* (**GatorDuo**_*α*_) reduces adaptability to dataset-specific domain granularity. This is expected because the KMeans-based refinement is highly sensitive to *K*: overly small *K* may merge distinct regions, while overly large *K* can fragment coherent domains and yield unstable pruning. Omitting MoE fusion or its associated loss (**GatorDuo**_*δ*_/**GatorDuo**_*γ*_) limits **GatorDuo**’s ability to effectively integrate complementary signals from the two views. This suggests that both the fusion mechanism and its auxiliary objective contribute to robust spatial domain identification.

**Fig. 3.**
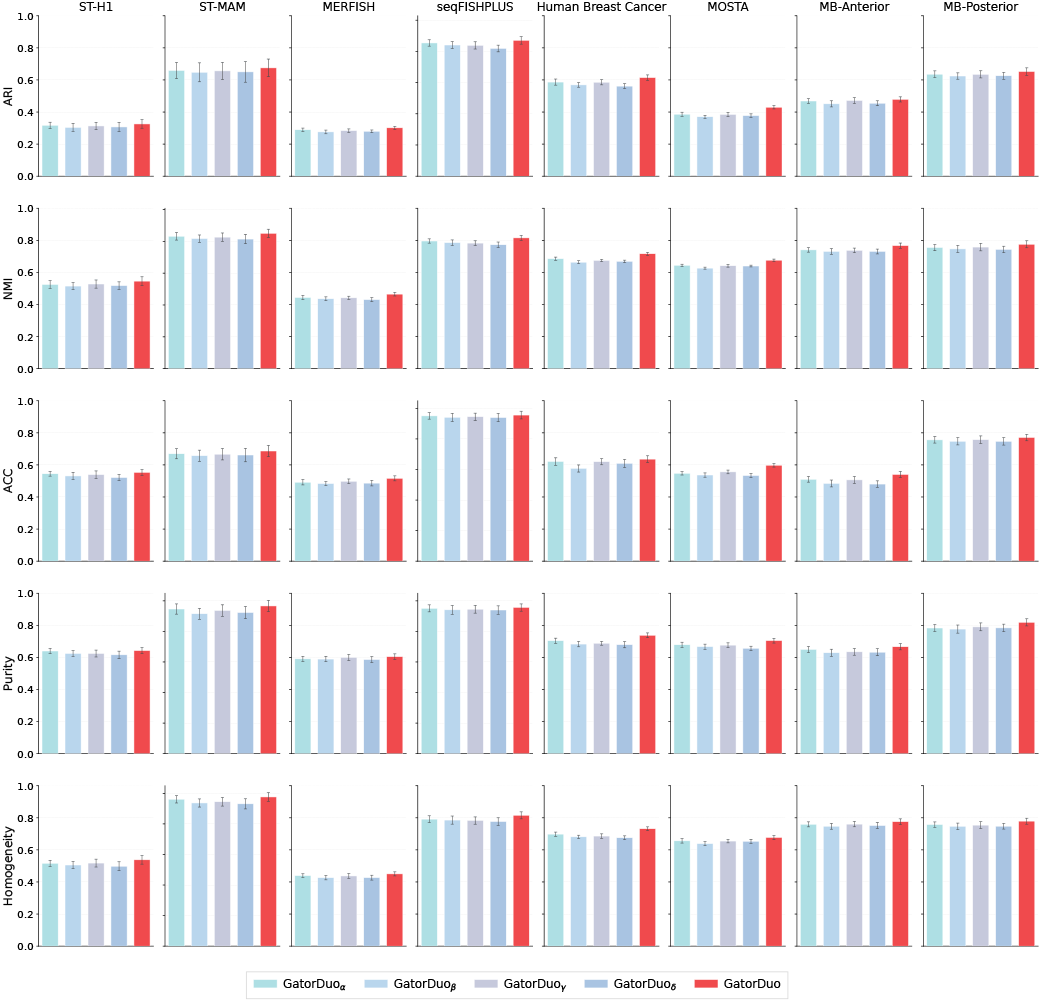
Performance of the full model and four ablation variants across the same eight datasets and five metrics. GatorDuo*α* (w/o RL): fixed *K* without RL-based selection; GatorDuo_*β*_ (w/o update-graph): removing the graph update/refinement module; GatorDuo*γ* (w/o MoE Loss): retaining MoE fusion but removing the MoE-specific loss term; GatorDuo_*δ*_ (w/o MoE): removing MoE and fusing two views by simple averaging.

### C. Hyperparameter sensitivity analysis

We further assess the robustness of **GatorDuo** to key hyperparameters that regulate (i) contrastive self-supervision, (ii) network regularization, and (iii) the relative contributions of major training signals. Specifically, we analyze the sensitivity to the temperature parameter *τ* used in contrastive self-supervision, the dropout rate used for the regularization of the neural-network, and the loss weighting coefficients (*λ*_CL_, *λ*_MOE_, *λ*_RL_) that balance the main loss components. We set *τ* to {0.1, 0.2, …, 0.8}, vary the dropout rate over {0.1, 0.2, 0.3, 0.4, 0.5}, and vary each of *λ*_CL_, *λ*_MOE_, and *λ*_RL_ over {0.1, 0.3, 0.5, 0.7, 0.9}. We evaluate spatial domain identification on eight benchmarks (ST-H1, ST-MAM, MERFISH, seqFISHPLUS, Human Breast Cancer, MOSTA, MB-Anterior, and MB-Posterior) using five clustering metrics: ARI, NMI, ACC, Purity, and Homogeneity. As shown in Figure 4, **GatorDuo** maintains stable performance across a broad range of *τ*, dropout rates, and loss-weight configurations across diverse datasets and evaluation criteria. Overall, these results indicate that **GatorDuo** is not overly sensitive to moderate variations in the contrastive temperature, regularization strength, or the loss trade-off coefficients, supporting the reliability of the reported gains under the default configuration.

**Fig. 4.**
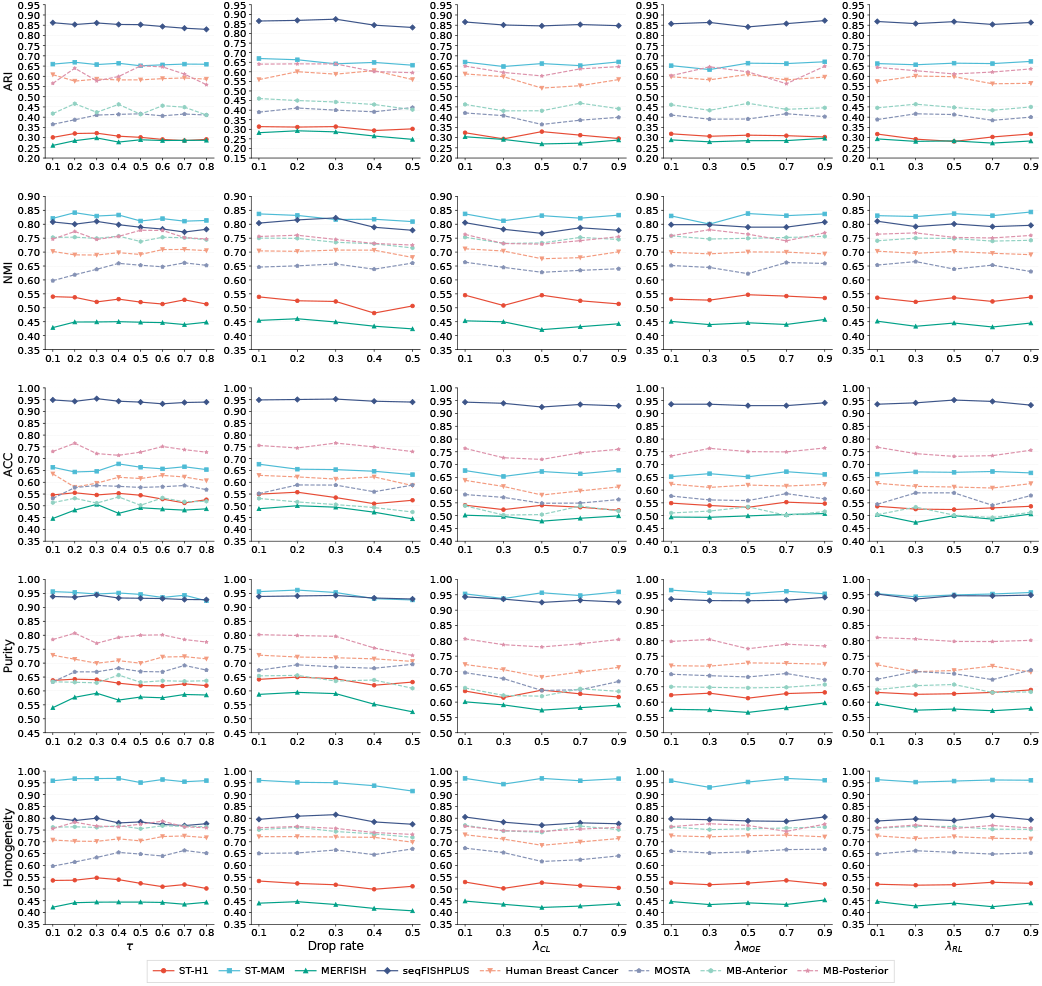
Hyperparameter sensitivity of GatorDuo for spatial domain identification across eight benchmarks. Rows report ARI, NMI, ACC, Purity, and Homogeneity, and columns correspond to the perturbed hyperparameter; each curve denotes one dataset.

### D. GatorDuo enables accurate spatial domain identification and coherent integration in mouse brain sections

We evaluated the spatial domain identification performance of **GatorDuo** on the two labeled sections, MB-Anterior and MB-Posterior, as shown in Figure 5. **GatorDuo** achieved the highest agreement with the ground-truth annotations in both sections, with an ARI of 0.441 on MB-Anterior and 0.557 on MB-Posterior, thereby outperforming all competing methods. On MB-Anterior, the strongest baseline methods were NichePCA (ARI = 0.396) and DMGCN (ARI = 0.382), whereas on MB-Posterior, spaVAE (ARI = 0.522) and BayesSpace (ARI = 0.511) showed the most competitive results. We next investigated the merged anterior-posterior sample, for which no ground-truth annotations were available. In this setting, performance was assessed primarily by qualitative evaluation, including spatial coherence, anatomical plausibility, and the ability to preserve biologically meaningful structure after section integration. In the merged sample, **GatorDuo** generated highly coherent spatial domains with clear laminar organization and minimal fragmentation. Furthermore, the learned topic proportion maps showed spatial patterns consistent with known cortical lamination (L2/3, L4, L5, L6, and L6b) and hippocampal subfields (CA1–CA3 and DG), suggesting that the integrated representation preserves biologically meaningful spatial organization.

**Fig. 5.**
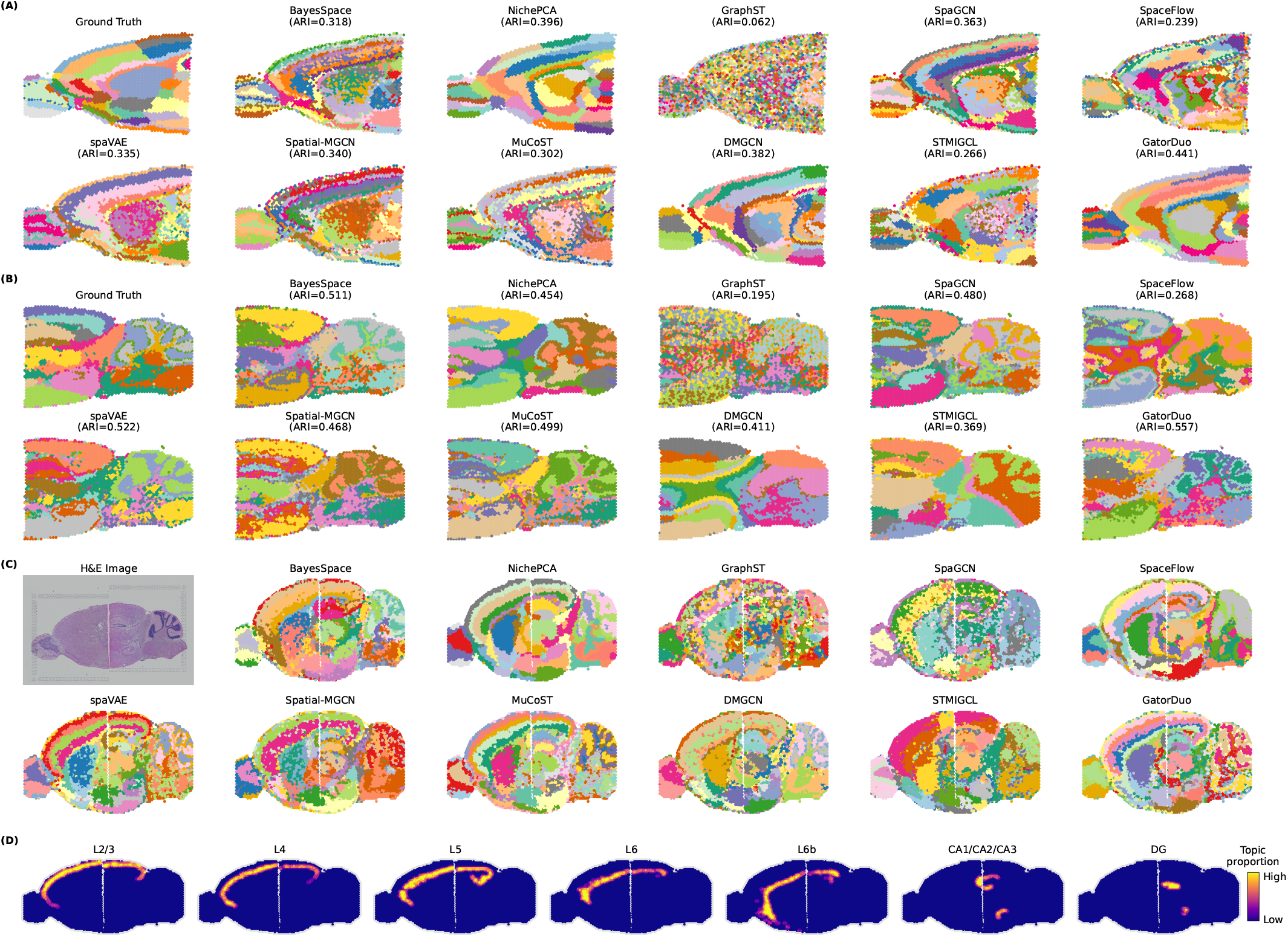
Spatial domain identification and section integration in the mouse brain. (A) Spatial domains in the labeled MB-Anterior section. (B) Spatial domains in the labeled MB-Posterior section. (C) Spatial domains in the merged anterior-posterior sample. (D) Topic proportion maps in the merged sample.

### E. GatorDuo recapitulates anatomical domains in the MOSTA E10.5 mouse embryo section

We evaluated **GatorDuo** on an annotated sagittal section from the MOSTA mouse embryo dataset at embryonic day 10.5 (E10.5; section E2S1) and compared its performance with ten representative spatial domain identification methods, as shown in Figure 6. Among all methods, **GatorDuo** achieved the highest concordance with the manual anatomical annotations, reaching an ARI of 0.398 and outperforming the strongest baseline, SpaGCN (ARI = 0.344). Beyond this quantitative improvement, **GatorDuo** generated spatial domains that were more contiguous and anatomically coherent, with sharper boundaries and closer agreement with the reference tissue map. The inferred domains recovered major embryonic structures, including the heart, liver, dermomyotome, lung primordium, and urogenital ridge. To further assess the biological validity of these domains, we examined the spatial expression of canonical marker genes. Consistent with the anatomical annotations, Myl7, Afp, Mylpf, Foxf1, and Wt1 were enriched in the heart, liver, dermomyotome, lung primordium, and urogenital ridge, respectively. Collectively, these results suggest that **GatorDuo** more accurately resolves both the spatial organization and biological identity of tissue domains in the developing mouse embryo.

**Fig. 6.**
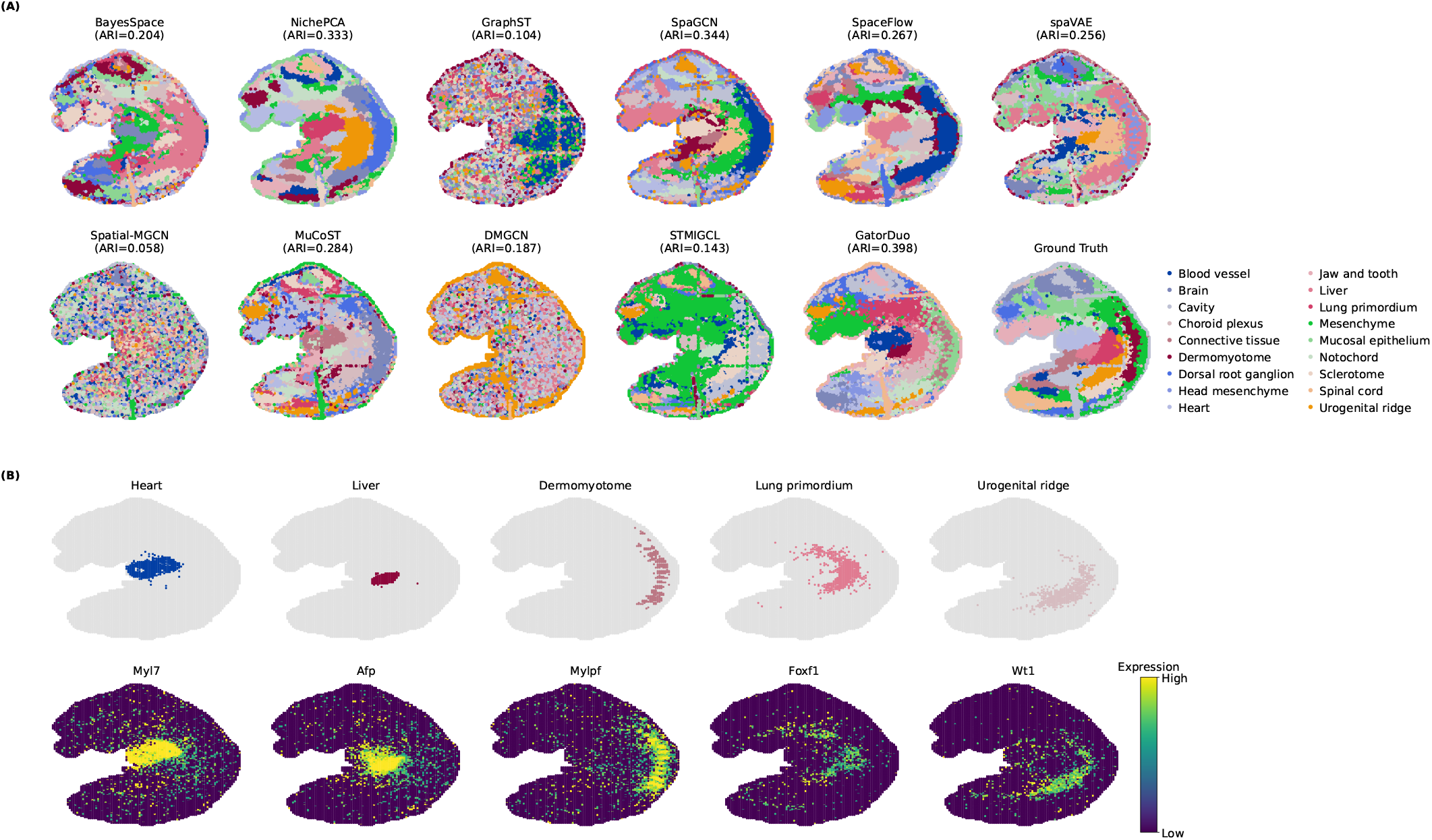
GatorDuo recapitulates anatomical domains in the MOSTA E10.5 mouse embryo section. (A) Spatial domain identification results generated by GatorDuo and ten baseline methods on an annotated sagittal section from the MOSTA mouse embryo dataset at embryonic day 10.5 (E10.5; section E2S1). (B) Spatial expression patterns of representative marker genes further support the biological relevance of the identified domains, including Myl7 for heart, Afp for liver, Mylpf for dermomyotome, Foxf1 for lung primordium, and Wt1 for urogenital ridge.

### F. GatorDuo accurately recovers histopathological architecture and reveals biologically meaningful spatial programs in human breast cancer

To assess the ability of spatial transcriptomics methods to recover histopathological architecture in a human breast cancer tissue section, we benchmarked multiple representative approaches against the annotated ground truth, as shown in Figure 7. Among all methods, **GatorDuo** achieved the highest agreement with manual annotations, with an ARI of 0.669, outperforming NichePCA (0.588), STMIGCL (0.492), SpaGCN (0.443), and BayesSpace (0.434). To further evaluate the biological relevance of the spatial domains identified by **GatorDuo**, we performed DisGeNET-based disease enrichment and TRRUST-based transcriptional regulator enrichment analyses on the genes associated with the identified regions. DisGeNET enrichment analysis [32] revealed significant enrichment of Breast Carcinoma, together with multiple mitochondria- and metabolism-related disease terms, suggesting that the genes underlying the recovered spatial domains are linked to molecular programs consistent with mitochondrial dysfunction and metabolic rewiring in breast cancer. TRRUST analysis [33] further identified several top-ranked transcriptional regulators, including RB1, E2F1, E2F3, MYBL2, NRF1, HIF1A, ATF3, and TP73, implicating regulatory programs involved in cell-cycle progression, mitochondrial and metabolic regulation, hypoxia-responsive signaling, and stress-adaptive transcription in breast cancer. These results suggest that the spatial domains recovered by **GatorDuo** are not only highly consistent with histopathological annotations but also supported by biologically meaningful disease associations and regulatory programs relevant to breast cancer.

**Fig. 7.**
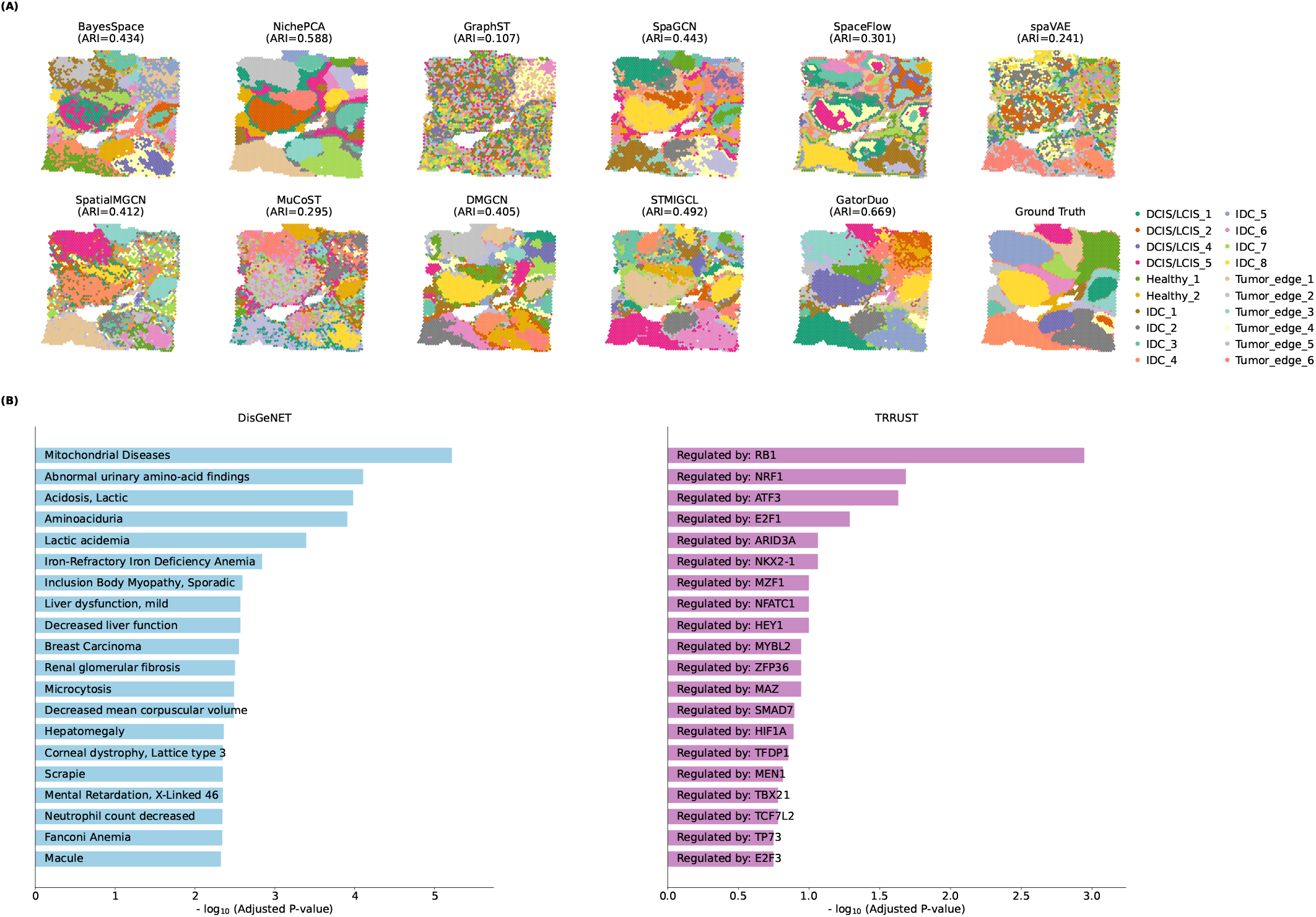
(A) Comparison of spatial domain identification results generated by different spatial transcriptomics methods on a human breast cancer section, together with the annotated ground truth. (B) Biological interpretation of genes associated with the spatial domains identified by GatorDuo.

## V. Conclusion

This work presents topology-aware self-supervised learning with explicit neighborhood correction as an effective framework for spatial domain identification in spatial transcriptomics. The proposed **GatorDuo** demonstrates that explicitly regularizing graph topology during self-supervised representation learning can strengthen spatial domain identification in spatial transcriptomics. By combining (i) pseudo-label agreement global-consistency graph refinement to suppress inconsistent cross-domain connections, (ii) contextual-bandit reinforcement learning to automatically select the refinement granularity (i.e., the number of clusters), and (iii) hybrid-routing MoE fusion to integrate transcriptomic-similarity and spatial-proximity views, the framework yields discriminative embeddings for downstream clustering and can further support downstream biological analyses. Across eight public benchmarks covering both sequencing- and imaging-based platforms and evaluated against ten competitive state-of-the-art baselines, **GatorDuo** achieves consistently strong performance under multiple clustering metrics. Ablation and sensitivity analyses further suggest that each component yields complementary gains in accuracy and robustness. Future work will generalize **GatorDuo** from 2D single-section analyses to serial-section and reconstructed 3D settings by leveraging inter-slice alignment and cross slice graph links to enforce more consistent domain identification across slices.

## Footnotes

1 https://www.10xgenomics.com/datasets/human-breast-cancer-block-a-section-1-1-standard-1-1-0

2 https://www.10xgenomics.com/datasets/mouse-brain-serial-section-2-sagittal-anterior-1-standard

3 https://www.10xgenomics.com/datasets/mouse-brain-serial-section-2-sagittal-posterior-1-standard

## REFERENCES

[1] A. Andersson, L. Larsson, L. Stenbeck, F. Salmén, A. Ehinger, S. Z. Wu, G. Al-Eryani, D. Roden, A. Swarbrick, Å. Borg et al., “Spatial deconvolution of her2-positive breast cancer delineates tumor-associated cell type interactions,” Nature communications, vol. 12, no. 1, p. 6012, 2021.

[2] C. Ortiz, J. F. Navarro, A. Jurek, A. Märtin, J. Lundeberg, and K. Meletis, “Molecular atlas of the adult mouse brain,” Science advances, vol. 6, no. 26, p. eabb3446, 2020.

[3] C. Xia, H. P. Babcock, J. R. Moffitt, and X. Zhuang, “Multiplexed detection of rna using merfish and branched dna amplification,” Scientific reports, vol. 9, no. 1, p. 7721, 2019.

[4] C.-H. L. Eng, M. Lawson, Q. Zhu, R. Dries, N. Koulena, Y. Takei, J. Yun, C. Cronin, C. Karp, G.-C. Yuan et al., “Transcriptome-scale super-resolved imaging in tissues by rna seqfish+,” Nature, vol. 568, no. 7751, pp. 235–239, 2019.

[5] A. Chen, S. Liao, M. Cheng, K. Ma, L. Wu, Y. Lai, X. Qiu, J. Yang, J. Xu, S. Hao et al., “Spatiotemporal transcriptomic atlas of mouse organogenesis using dna nanoball-patterned arrays,” Cell, vol. 185, no. 10, pp. 1777–1792, 2022.

[6] Z. Wang, A. Geng, H. Duan, F. Cui, Q. Zou, and Z. Zhang, “A comprehensive review of approaches for spatial domain recognition of spatial transcriptomes,” Briefings in Functional Genomics, vol. 23, no. 6, pp. 702–712, 2024.

[7] S. Wang, Y. Liu, Z. Zhang, Q. Song, and J. Bian, “Gatorst: A versatile contrastive meta-learning framework for spatial transcriptomic data analysis,” bioRxiv, 2025.

[8] L. Zhang, S. Liang, and L. Wan, “A multi-view graph contrastive learning framework for deciphering spatially resolved transcriptomics data,” Briefings in Bioinformatics, vol. 25, no. 4, p. bbae255, 2024.

[9] E. Zhao, M. R. Stone, X. Ren, J. Guenthoer, K. S. Smythe, T. Pulliam, S. R. Williams, C. R. Uytingco, S. E. Taylor, P. Nghiem et al., “Spatial transcriptomics at subspot resolution with bayesspace,” Nature biotechnology, vol. 39, no. 11, pp. 1375–1384, 2021.

[10] D. P. Schaub, B. Yousefi, N. Kaiser, R. Khatri, V. G. Puelles, C. F. Krebs, U. Panzer, and S. Bonn, “Pca-based spatial domain identification with state-of-the-art performance,” Bioinformatics, vol. 41, no. 1, p. btaf005, 2025.

[11] Y. Long, K. S. Ang, M. Li, K. L. K. Chong, R. Sethi, C. Zhong, H. Xu, Z. Ong, K. Sachaphibulkij, A. Chen et al., “Spatially informed clustering, integration, and deconvolution of spatial transcriptomics with graphst,” Nature Communications, vol. 14, no. 1, p. 1155, 2023.

[12] J. Hu, X. Li, K. Coleman, A. Schroeder, N. Ma, D. J. Irwin, E. B. Lee, R. T. Shinohara, and M. Li, “Spagcn: Integrating gene expression, spatial location and histology to identify spatial domains and spatially variable genes by graph convolutional network,” Nature methods, vol. 18, no. 11, pp. 1342–1351, 2021.

[13] H. Ren, B. L. Walker, Z. Cang, and Q. Nie, “Identifying multicellular spatiotemporal organization of cells with spaceflow,” Nature communications, vol. 13, no. 1, p. 4076, 2022.

[14] T. Tian, J. Zhang, X. Lin, Z. Wei, and H. Hakonarson, “Dependency-aware deep generative models for multitasking analysis of spatial omics data,” Nature Methods, vol. 21, no. 8, pp. 1501–1513, 2024.

[15] B. Wang, J. Luo, Y. Liu, W. Shi, Z. Xiong, C. Shen, and Y. Long, “Spatial-mgcn: a novel multi-view graph convolutional network for identifying spatial domains with attention mechanism,” Briefings in Bioinformatics, vol. 24, no. 5, p. bbad262, 2023.

[16] X. Shi, J. Zhu, Y. Long, and C. Liang, “Identifying spatial domains of spatially resolved transcriptomics via multi-view graph convolutional networks,” Briefings in Bioinformatics, vol. 24, no. 5, 2023.

[17] X. Liang, S. Xiao, L. Ba, Y. Feng, Z. Ma, F. Adilova, J. Qi, and S. Jin, “Spatial domain identification method based on multi-view graph convolutional network and contrastive learning,” PLOS Computational Biology, vol. 21, no. 10, p. e1013369, 2025.

[18] S. Ren, X. Liao, F. Liu, J. Li, X. Gao, and B. Yu, “Exploring the latent information in spatial transcriptomics data via multi-view graph convolutional network based on implicit contrastive learning,” Advanced Science, vol. 12, no. 21, p. 2413545, 2025.

[19] X. Wang, S. Wang, X. Liang, D. Zhao, J. Huang, X. Xu, B. Dai, and Q. Miao, “Deep reinforcement learning: A survey,” IEEE Transactions on Neural Networks and Learning Systems, vol. 35, no. 4, pp. 5064–5078, 2022.

[20] A. Agarwal, S. M. Kakade, J. D. Lee, and G. Mahajan, “On the theory of policy gradient methods: Optimality, approximation, and distribution shift,” Journal of Machine Learning Research, vol. 22, no. 98, pp. 1–76, 2021.

[21] A. Bietti, A. Agarwal, and J. Langford, “A contextual bandit bake-off,” Journal of Machine Learning Research, vol. 22, no. 133, pp. 1–49, 2021.

[22] R. J. Williams, “Simple statistical gradient-following algorithms for connectionist reinforcement learning,” Machine learning, vol. 8, no. 3, pp. 229–256, 1992.

[23] J. Zhang, J. Kim, B. O’Donoghue, and S. Boyd, “Sample efficient reinforcement learning with reinforce,” in Proceedings of the AAAI conference on artificial intelligence, vol. 35, no. 12, 2021, pp. 10887–10895.

[24] R. A. Jacobs, M. I. Jordan, S. J. Nowlan, and G. E. Hinton, “Adaptive mixtures of local experts,” Neural computation, vol. 3, no. 1, pp. 79–87, 1991.

[25] N. Shazeer, A. Mirhoseini, K. Maziarz, A. Davis, Q. Le, G. Hinton, and J. Dean, “Outrageously large neural networks: The sparsely-gated mixture-of-experts layer,” arXiv preprint arXiv:1701.06538, 2017.

[26] W. Fedus, B. Zoph, and N. Shazeer, “Switch transformers: Scaling to trillion parameter models with simple and efficient sparsity,” Journal of Machine Learning Research, vol. 23, no. 120, pp. 1–39, 2022.

[27] T. Kipf, “Semi-supervised classification with graph convolutional networks,” arXiv preprint arXiv:1609.02907, 2016.

[28] Z. Zhang, Y. Liu, M. Xiao, K. Wang, Y. Huang, J. Bian, R. Yang, and F. Li, “Graph contrastive learning as a versatile foundation for advanced scrna-seq data analysis,” Briefings in Bioinformatics, vol. 25, no. 6, p. bbae558, 2024.

[29] Z. Zhang, Y. Liu, J. Bian, A. Jimeno Yepes, J. Shen, F. Li, G. Long, and F. D. Salim, “Boosting patient representation learning via graph contrastive learning,” in Joint European Conference on Machine Learning and Knowledge Discovery in Databases. Springer, 2024, pp. 335–350.

[30] Y. Liu, Z. Zhang, J. Mi, S. Pan, T. Chen, Y. Guo, X. He, and J. Bian, “Gatorclr: Personalized predictions of patient outcomes on electronic health records using self-supervised contrastive graph representation,” Journal of Biomedical Informatics, p. 104851, 2025.

[31] A. van den Oord, Y. Li, and O. Vinyals, “Representation learning with contrastive predictive coding,” arXiv preprint arXiv:1807.03748, 2018.

[32] J. Piñero, N. Queralt-Rosinach, A. Bravo, J. Deu-Pons, A. Bauer-Mehren, M. Baron, F. Sanz, and L. I. Furlong, “Disgenet: a discovery platform for the dynamical exploration of human diseases and their genes,” Database, vol. 2015, p. bav028, 2015.

[33] H. Han, H. Shim, D. Shin, J. E. Shim, Y. Ko, J. Shin, H. Kim, A. Cho, E. Kim, T. Lee et al., “Trrust: a reference database of human transcriptional regulatory interactions,” Scientific reports, vol. 5, no. 1, p. 11432, 2015.

